# DNA as a Recyclable Natural Polymer

**DOI:** 10.1101/2021.08.31.458327

**Authors:** Weina Liu, Simone Giaveri, Daniel Ortiz, Francesco Stellacci

## Abstract

Nature has the ability of circularly re-using its components to produce molecules and materials it needs. An example is the ability of most living organisms of digesting proteins they feed off into amino acids and then using such amino acids in the ribosomal synthesis of new proteins. Recently, we have shown that such recycling of proteins can be reproduced outside living organisms. The key proteins’ feature that allows for this type of recycling is their being sequence-defined polymers. Arguably, Nature’s most famous sequence-defined polymer is DNA. Here we show that it is possible starting from sheared calf-DNA to obtain all the four nucleotides as monophosphate-nucleotides (dNMPs). These dNMPs were phosphorylated in a one-pot, multi-enzymes, phosphorylation reaction to generate triphosphate-nucleotides (dNTPs). Finally, we used the dNTPs so achieved (with a global yield of ∼60%) as reagents for PCR (polymerase chain reaction) to produce target DNA strands, and for the diagnose of targeted DNA by quantitative PCR (qPCR). This approach is an efficient, convenient, and environmentally friendly way to produce dNTPs and DNA through recycling according to the paradigm of circular economy.

## 1. Introduction

Nature can efficiently break down complex biomass into small molecules and circularly use them to fulfill its requirement of biosynthesis, a recycling masterpiece. For example, bacteria in stomach-intestinal system can efficiently digest food to obtain nutrition molecules for the synthesis of bioactive macromolecules.^[1]^ Also, microorganisms used in fermentation processes can degrade organic nutrients to alcohol or other small molecules. Those functions have been widely utilized in alcoholic beverages or dairy industry,^[2,3]^ while most other approaches that Nature has to ‘recycle’ proteins and nucleic acids have not yet been translated in laboratory settings. Recently we have shown that it is possible to reproduce outside living organisms the approach organisms use to ‘recycle’ mixtures of n proteins into the (n+1)^th^ protein of interest, not necessarily related to the parent ones.Briefly, protein mixtures were first enzymatically digested into their constitutive amino acids, and then a cell-free transcription-translation system was used to recycle the so obtained amino acids into fluorescent (GFP, mScarlet-i), or bioactive proteins (catechol 2,3-dioxygenase), by means of the ribosomal expression. Recycling unseparated mixtures of proteins into the protein of need, not necessarily related to the parent materials, can only work for sequence-defined polymers, where function derives from the sequence of monomers.

DNA is also a sequence-defined polymer where function derived from the exact order of the four nucleotides (bases) that determine its sequence. Hence, following the same principles illustrated for the proteins/amino acids system, one can conceive an approach where a random mixture of DNA can be ‘digested’ into its nucleotides that can then be put back together into a new DNA sequence unrelated to the original one. In the case of DNA the formation of the final DNA product can be made with a man-invented process (polymerase chain reaction, PCR).^[5]^ When compared to protein recycling, DNA recycling requires an extra step.^[6]^ DNA can be depolymerized through an enzymatically hydrolysis step to break inter-nucleotide phosphodiester backbone, but the product of such reaction are monophosphate-nucleotides (dNMPs). Unfortunately, dNMPs cannot be directly used as reagents to produce new DNA sequences, one needs to first convert them into triphosphate-nucleotides (dNTPs) by introducing additional phosphate groups. In this work, we used Nature available DNA (calf DNA) as a starting material to mimic the Nature’s recycling of DNA. We first established a method to depolymerize DNA into dNMPs with efficiency exceeding 80%. We then developed a convenient, efficient synthesis to phosphorylate the mixture of the four dNMPs into the respective dNTPs with yields more than 95% (commercialized dNMPs) and ∼80% (recycled dNMPs). Finally, we proved that the so produced dNTPs could be used for the biosynthesis of new DNA sequences by PCR amplification, and for nucleic acid testing by quantitative PCR (qPCR). This methodology could bring a totally new path to obtain monomeric nucleotide materials, which might replace their chemical synthesis.^[7]^

## 2. Result and discussion

### 2.1. DNA De-polymerization

Nature is a vast storage of polymeric DNA materials, e.g., the maximum DNA content can be 10% (dry mass basis) from bacterioplankton.^[8]^ Also, Nature can rapidly and constantly produce DNA, which can be considered as a sustainable resource to recycle monomeric nucleotide materials. In this work, commercially available calf thymus DNA (calf DNA) with known content of CG 41.9% and AT 58.1% (specifications from Sigma-Aldrich) was used as a natural DNA source for the enzymatic hydrolysis to obtain nucleotide monomers (**Figure 1a**). To efficiently hydrolyse calf DNA and release dNMPs, it is important to choose the cleavage site at 3’-terminal of the phosphodiester bonds, so that the monophosphate group can be kept at 5’-terminal of the released monomeric nucleotides. To obtain the desired hydrolysis product, we chose Exonuclease III (Exo III) and Exonuclease I (Exo I) with the required 3’-phosphomonoesterase activity. Exo III is a double-strand DNA (dsDNA) specific exonuclease, which can catalyze the stepwise removal of dNMPs from 3’-terminal of dsDNA with blunt end or 5’-overhang.^[9,10]^ Exo I is a single-strand DNA (ssDNA) specific exonuclease, which can catalyze the stepwise removal of dNMPs from ssDNA in the 3’- to 5’-direction.^[11]^ The mixture of calf DNA, Exo III, and Exo I was incubated in 1x Exo III buffer at 37°C overnight for hydrolysis (Figure 1a, step 1). Afterwards, unhydrolyzed, and hydrolyzed calf DNA were loaded to a 2% agarose gel. In Figure 1b we show a representative image of a gel containing the starting materials as well as the hydrolyzed one. The initial calf DNA shows a smeared band (lane 2, sequence length between 100-2000 and above 2000 base pair), the hydrolyzed one has an almost absent band (lane 3), indicating the relatively high hydrolysis efficiency.

**Figure 1.**
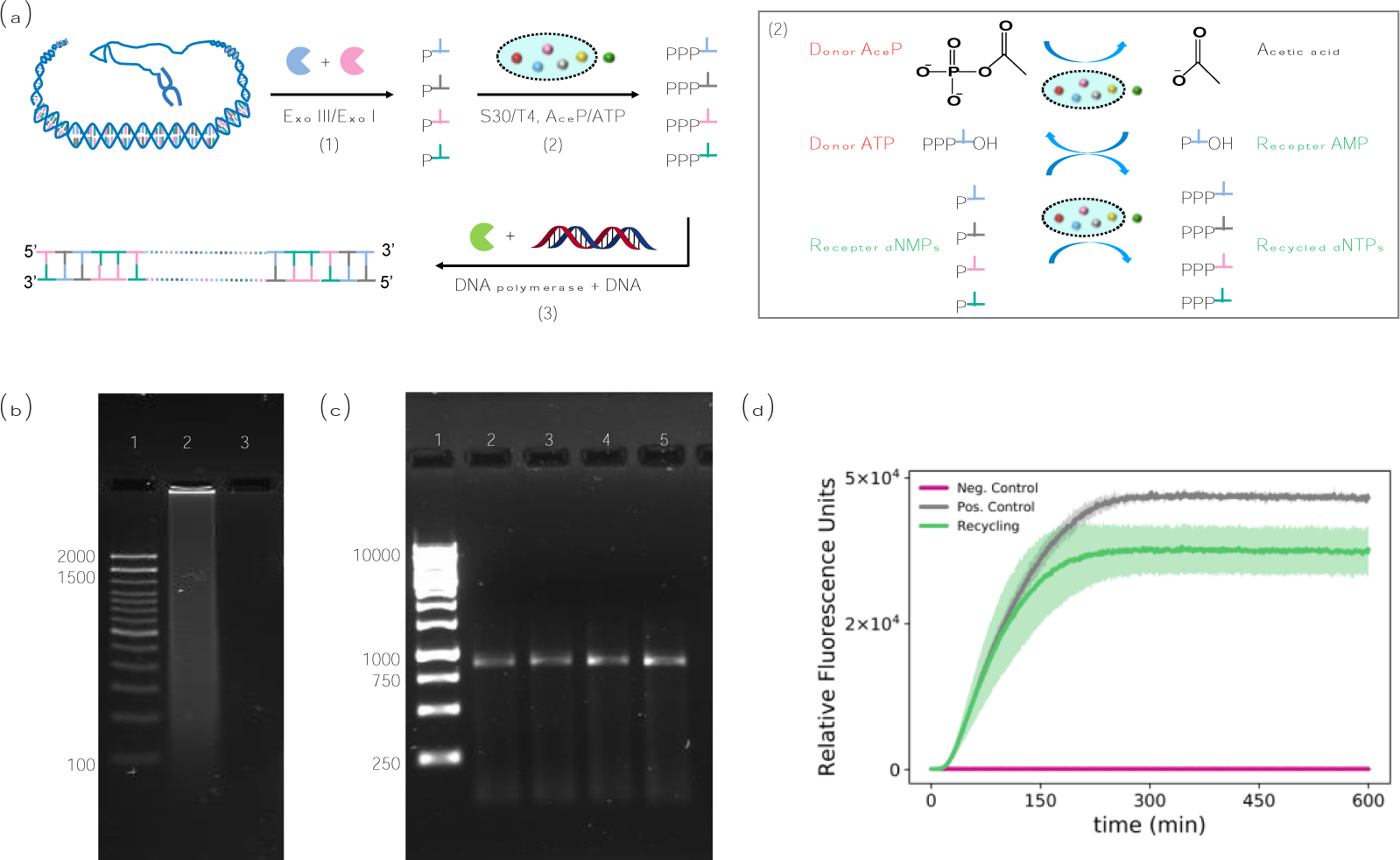
DNA recycling from Nature. (a) Scheme of monomers recycling from calf thymus DNA for PCR, step (1) calf DNA hydrolysis by Exo III and Exo I to release dNMPs, step (2) dNMPs_calf DNA phosphorylation, step (3) PCR to amplify new DNA sequences. The step (2) phosphorylation reaction (insert) was a one-pot reaction catalyzed by mixture of E. coli S30 extract, as E. coli S30 extract was found rich in acetyl kinase^[41]^ and nucleotide mono-/di-phosphate kinase,^[24,42]^ and T4 NMP Kinase can catalyze the phosphorylation of dCMP, dTMP and dGMP. (b) Agarose gel (2%) of ladder (lane 1), calf DNA (lane 2), hydrolyzed calf DNA (lane 3). (c) Agarose gel (1%) of the PCR amplified GFP sequence from recycled dNTPs (duplicate, lane 2 and 3), as well as from the purchased dNTPs as positive control (duplicate, lane 4 and 5). (d) Plots of the fluorescence signal resulting from the expression of GFP in the TX-TL system. The green curve is obtained by feeding the TX-TL system with a GFP DNA template (75 ng) polymerized from the recycled dNTPs. The grey curve (positive control) is obtained as the result of an expression experiment with the TX-TL system supplemented with a GFP DNA template (75 ng) polymerized from purchased dNTPs. In the negative control expression (violet curve), the TX-TL system was not supplemented with any DNA template.

To identify the DNA hydrolysis product, and to determine the hydrolysis efficiency, we used LC-MS for qualitative as well as quantitative analysis. The calibration curves from dNMPs standard solutions can be found in supporting information (Figure S1a). The plot of XIC (extracted-ion chromatogram) of the hydrolysis product is shown in Figure 2a. It illustrates that the mixture of Exo III and Exo I could efficiently hydrolyze DNA to release dNMPs. The average hydrolysis efficiency of four dNMPs was 83.9 ± 0.6% (Figure S1b). The possible reason of non-completely hydrolysis could be the hydrolysis efficacy of nuclease Exo III, which can hydrolyze dsDNA with blunt end and 5’-overhang but cannot hydrolyze dsDNA with 3’-overhang. As the calf DNA was mechanically sheared, it was not possible to exclude the existence of dsDNA with 3’-overhang in the starting material of the DNA hydrolysis reaction. Overall, the calf DNA hydrolysis was relatively efficient with desired product and high yield. The established hydrolysis method is non-selective for sequence; hence we believe that it will be applicable to all kinds of sheared DNA.

**Figure 2.**
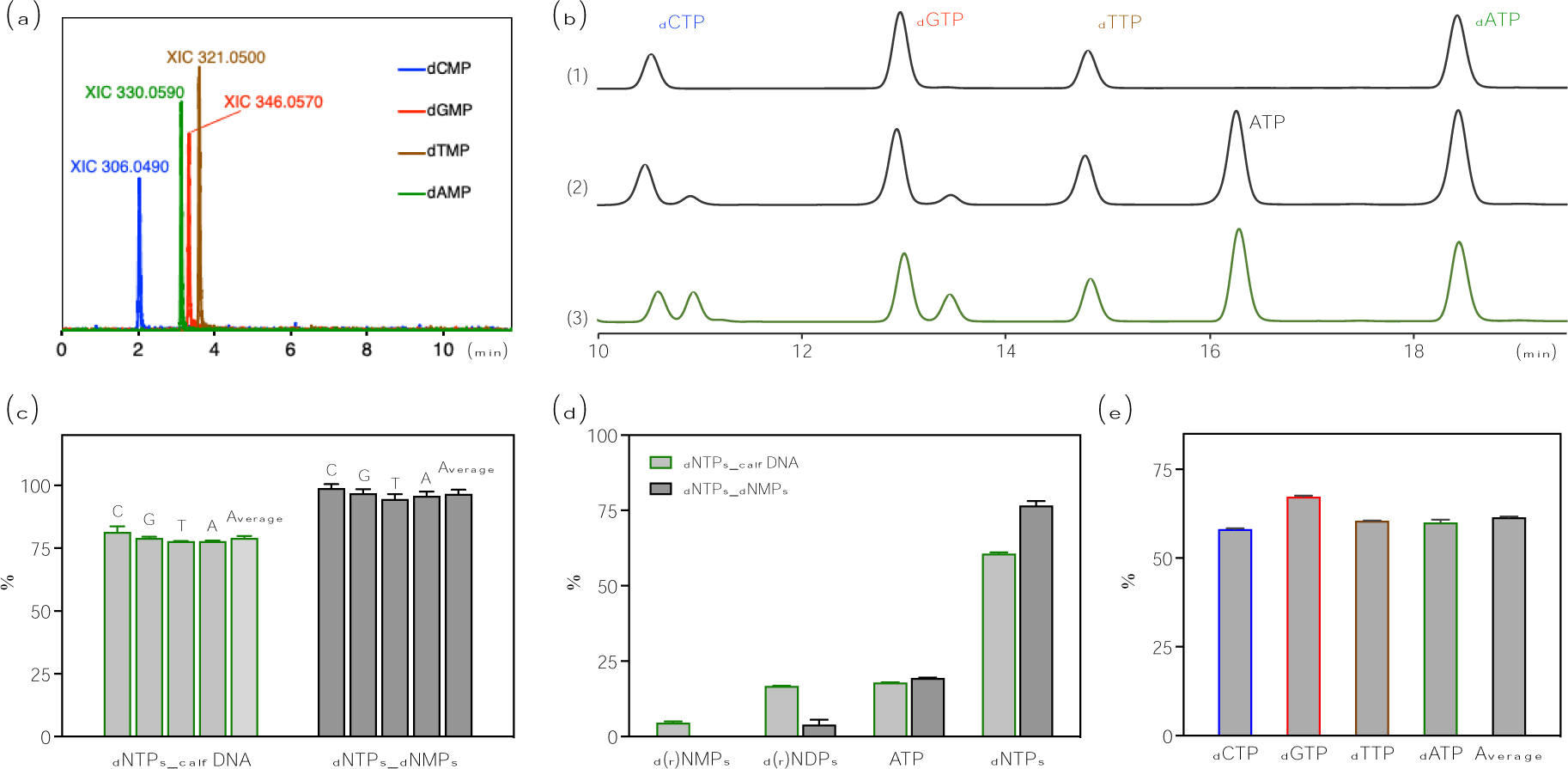
Quantification of calf DNA hydrolysis and dNMPs phosphorylation efficiency by HPLC and HPLC-MS. (a) Extracted-ion chromatogram (XIC) from LC-MS analysis of dNMPs_calf DNA (dAMP_330.0590, dCMP_306.0490, dTMP_321.0500, dGMP_346.0570). (b)HPLC retention time of (1) dNTPs standards, (2) dNTPs_dNMPs, and (3) dNTPs_calf DNA with spectra of full retention time in Figure S2. (c) Phosphorylation efficiency of dNTPs_calf DNA (C 81.4 ± 2.2%, G 79.1 ± 0.4%, T 77.7 ± 0.1%, A 77.8 ± 0.2%, and in average 79.0 ± 0.7%), and phosphorylation efficiency of dNTPs_dNMPs (C 98.9 ± 1.5%, G 96.8 ± 1.6%, T 94.5 ± 1.9%, A 95.9 ± 1.5%, and in average 96.5 ± 1.6%). (d) Percentage of all the phosphorylation products and residues from the phosphorylation reaction of dNTPs_calf DNA (d(r)NMPs (dNMPs and AMP (ATP hydrolysis product)) 4.6 ± 0.4%%, d(r)NDPs (dNDPs and ADP (ATP hydrolysis products)) 16.7 ± 0.1%, ATP 17.9 ± 0.1%, dNTPs 60.6 ± 0.3%); and percentage of all the phosphorylation products and residues from the phosphorylation reaction of dNTPs_dNMPs (d(r)NMPs 0 %, d(r)NDPs 3.9 ± 1.6%, ATP 19.4 ± 0.1%, dNTPs 76.6 ± 1.4%). Calibration curves of d(r)NDPs and d(r)NTPs can be found in Figure S3a and S3b. Concentration of dNDPs and dNTPs in the phosphorylation product can be found in Figure S3c and S3d. The quantification of d(r)NMPs was obtained by area integration due to very low residue. (e) Recycling efficiency of dNTPs from calf DNA (dCTP 58.1 ± 0.24%, dGTP 67.2 ± 0.2 %, dTTP 60.5 ± 0.03%, dATP 60.0 ± 0.8%, in average 61.5 ± 0.2%).

### 2.2. One-pot dNMPs phosphorylation

As previously mentioned, the DNA hydrolysis products dNMPs could not be directly applied for the biosynthesis of new DNA sequence, as this required dNTPs. To address this problem, we implemented a phosphorylation step to convert the dNMPs into dNTPs. Since the DNA hydrolysis product was a mixture of four dNMPs, it was important to establish a phosphorylation approach that could convert the four dNMPs into dNTPs in a one-pot reaction. We looked for a multi-enzymes bio-catalysis as an efficient approach.^[12,13]^ In the biosystem, the phosphorylation of nucleotides is catalyzed by intracellular phosphotransferase and kinase.^[14,15]^ Therefore, cell lysate is an excellent catalyst and has been applied for this nucleotide phosphorylation reaction,^[16,17]^ but the reported phosphorylation yields are relatively low (especially for dTTP). We thought that a possible reason for such low yields could be the imbalanced content of four nucleotides in the intracellular nucleotide pool. It is known that such pool is rich in A and G bases as they are also needed for energy storage, signaling and apoptosis.^[18]^ Hence one can postulate that enzymes to phosphorylate the T and C bases less present than the ones to perform the same task in the other bases.^[19]^ To improve the phosphorylation yield of this one-pot reaction, we added T4-nucleotide monophosphate Kinase (T4), that can specifically catalyze the phosphorylation of dTMP, dCMP, and dGMP.^[20]^ We performed the one-pot phosphorylation in an acetyl-phosphate/ATP dual-phosphate-donors system (Figure 1a, step 2, details see insert), with the mixture of E. coli S30 cell extract (S30) and T4 as catalyst. S30 is a commercial cell lysate extract product,^[21–23]^ that has been originally established for cell-free protein expression.^[24]^ ATP was used as a phosphate donor and a cofactor, that was continuously consumed and re-generated, see insert in Figure 1a. Acetyl-phosphate (AceP) was applied as the phosphate donor to re-generate ATP. During this phosphorylation step, a phosphate group was introduced to dNMPs, with dNDPs (nucleotide diphosphate) formed as intermediate products, and further a second phosphate group was introduced to dNDPs to generate the mixture of dNTPs.

To evaluate the phosphorylation efficiency of this one-pot reaction, we first performed the reaction with a commercial dNMPs mixtures with equal content of four bases. The enzyme mixture was carefully adjusted to achieve the best phosphorylation efficiency of all four dNMPs. The phosphorylation product referred as dNTPs_dNMPs was quantified by HPLC^[25]^ (retention time of the dNTPs products in Figure 2b, line 2; full retention time of all components in Figure S2, line 4). After the one-pot phosphorylation reaction, the dNMPs residues were not detectable anymore, and the amount of intermediate phosphorylation product dNDPs was quite low. A relatively high average phosphorylation yield was achieved (96.5 ± 1.6%, Figure 2c), which is higher than the reported nucleotide phosphorylation by chemistry methods.^[7,26,27]^ In addition to the relatively good phosphorylation efficiency, this one-pot enzymatic reaction has many other advantages, such as mild reaction condition, green chemistry, all aqueous medium without any organic solvent, one-pot reaction condition suitable for multiple phosphate-receptors (four dNMPs and four dNDPs). In the phosphorylation mixture, final content of dNTPs was 76.6 ± 1.4%, the d(r)NMPs residue was 0%, d(r)NDPs residue was 3.9 ± 1.6% (with the concentration of each dNDPs in Figure S3c), and ATP residue was 19.4 ± 0.1% (Figure 2d).

Further, phosphorylation of recycled dNMPs_calf DNA was performed by the same approach, and the product referred as dNTPs_calf DNA was quantified by HPLC (chromatography of the dNTPs products in Figure 2b, line 3; full chromatography of all components in Figure S2, line 5). The average phosphorylation efficiency of dNTPs recycled from calf DNA was calculated to be 79.0 ± 0.7% (Figure 2c). There was 18.1% decreased in phosphorylation efficiency of dNTPs_calf DNA in comparison to the dNTPs_dNMPs standards. The possible reason could be the unequal ratio of four nucleobases from recycled dNMPs, which led to unequal catalysis efficiency in this one-pot, competing phosphorylation reaction. Nevertheless, the phosphorylation efficiency is still relatively good. In the phosphorylation mixture, dNTPs_calf DNA content was 60.6 ± 0.3%, d(r)NMPs residue was 4.6 ± 0.4%, d(r)NDPs residue was 16.7 ± 0.1% (with the concentration of each dNDPs in Figure S3c), and ATP residue was 17.9 ± 0.1% (Figure 2d). The concentration of all four dNTPs_calf DNA was in the range of 190-280 µM (in average 243.7 ± 0.9 µM, Figure S3d), which was suitable to be directly used for the synthesis of new DNA by PCR. Calculated from the total mass of starting material calf DNA, the average recycling efficiency of dNTPs_calf DNA was 61.5 ± 0.2% (all four dNTPs with recycling yield in the range of 58.1%-67.2%, Figure 2e), which was relatively good after two steps of hydrolysis and phosphorylation reactions.

### 2.3. DNA Re-polymerization

As a mimic of Nature DNA material circulation, we further re-polymerized the recycled dNTPs_calf DNA into a new DNA sequence by PCR (Figure 1a, step 3). In this work, we decided to use dNTPs_calf DNA directly for PCR after a simple filtration step without any further purification. We are aware that by doing so we did not remove ATP and dNDPs from dNTPs_calf DNA. Such impurities could potentially affect the efficiency of qPCR and also insert into the new sequence produced^[28]^. We judged that the level of such impurity was so low that an extra purification step was not justified. To test this hypothesis, we performed qPCR by adding 0.2 mM ATP to qPCR substrate that used commercial dNTPs and observed only a slight decrease in qPCR efficiency (Figure S5). We re-polymerized the recycled dNTPs by PCR amplification of a linear DNA template encoding GFP (Figure 1c, duplicate, lane 2 and 3). Commercially available dNTPs were used as positive control for PCR (Figure 1c, duplicate, lane 4 and 5). The so achieved DNA was purified, and fed into a commonly used cell-free transcription-translation (TX-TL) system^[29]^ (PUREfrex™, Kaneka Eurogentec SA, see Supporting Information) for verifying the “transcription”, and “translation” of the GFP sequence from a DNA template polymerized with recycled nucleotides. Upon feeding a DNA template, the protein of interest was expressed in the TX-TL system. As shown in Figure 1d, a good yield for GFP expression was achieved, proving that the re-polymerized DNA from the hydrolysed DNA precursor encodes a new genetic information for protein expression.

The GFP expression yield of recycled DNA was about 20% lower than positive control. There are many potential reasons for this, among them we can mention the insertion of 5mC (5-methylcytosine) into the new GFP sequence given that 5mC is an epigenetic factor for the regulation of gene expression^[30]^. In our case, 5mC could originate from dNMPs_calf DNA. Although 5mC was not detected from dNMPs_calf DNA by LC-MS, its natural abundance (about 0.88% in mammals’ genomic DNA^[31]^) cannot be neglected, and the possibility of circulating the 5mC from calf DNA into the new GFP sequence cannot be excluded. We tested by Nanopore sequencing the linear plasmid we used to express GFP and it showed a certain amount of 5mC. Further investigation must be performed to fully understand this issue.

### 2.4. Recycling DNA for qPCR

Quantitative PCR (qPCR) is a powerful molecular diagnose tool to quantify gene expression,^[33]^ which is largely consumed for nucleic acid testing during SARS-CoV-2 pandemic.^[34]^ Since Nature DNA can be efficiently recycled, next, we tested the possibility of using recycled dNTPs as substrate for qPCR. The cycles of threshold (C_T_) are defined as the qPCR cycle at which the fluorescent signal of the reporter dye crosses an arbitrarily placed threshold, indicating the exponential phase of qPCR amplification.^[35]^ The C_T_ inversely depends on DNA amplicon amount in the reaction mixture (the more DNA template, the lower the C_T_ value). With equal amount of amplicons, the C_T_ value as a reference of the qPCR performance is influenced by the other components, i.e., the concentration of dNTPs involved in this polymerization process,^[36– 38]^ and the content of impurities in PCR substrate. Therefore, in this work the performance of qPCR was used to evaluate the quality of recycled dNTPs.

Since the DNA recycling process is applicable to all genomic DNA, E. coli DNA as an alternative DNA resource was applied for recycling of monomeric dNTPs. The E. coli DNA was hydrolyzed, phosphorylated by the same protocol of calf DNA recycling (details see SI). The recycled product referred as dNTPs_E. coli DNA was also applied for nucleic acid testing by qPCR amplification. Self-made qPCR kits were prepared by mixture of DreamTaq polymerase, Sybr dye, recycled dNTPs (dNTPs_calf DNA or dNTPs_E. coli DNA). The average concentration of dNTPs was adjusted to 0.2 mM. Commercially available qPCR kit was applied as positive control. The self-made qPCR kits were used for the amplification of a fragment with 133 base pair from luciferase DNA template. Primers were designed by IDT PrimerQuest™ Tool. As shown in Figure 3a, similar C_T_ values of self-made qPCR kits and positive control were obtained with low content of DNA template (0.001 ng). With higher content of template DNA (1, 0.1, 0.01 ng), the detection performance of self-made qPCR kits was slightly lower than positive control with one or two more circles of C_T_ needed (Figure 3a, amplification plots see Figure S4). The potential effect of ATP and dNDPs residues to the qPCR performance was minimal (Figure S5). Since the commercially available qPCR kit (the positive control) is in fully optimized condition, the analytical performance of self-made qPCR kit from recycled dNTPs is still considered relatively good. For the no template control (NTC) sample, the C_T_ value of the commercially available qPCR kit was 35.3, The self-made qPCR assay was 30.8 and 30.4 (qPCR kit prepared from dNTPs_calf DNA and dNTPs_E. coli DNA), respectively. The detect limitation of self-made qPCR kits (template DNA content down to 0.001 ng) was not affected by NTC results. The qPCR amplification products were loaded to a Page-gel, showing that the amplification products with desired length (Figure 3b). As a molecular diagnosing tool, the preparation of self-made qPCR from recycled dNTPs is relatively convenient and its analytical performance is very good.

**Figure 3.**
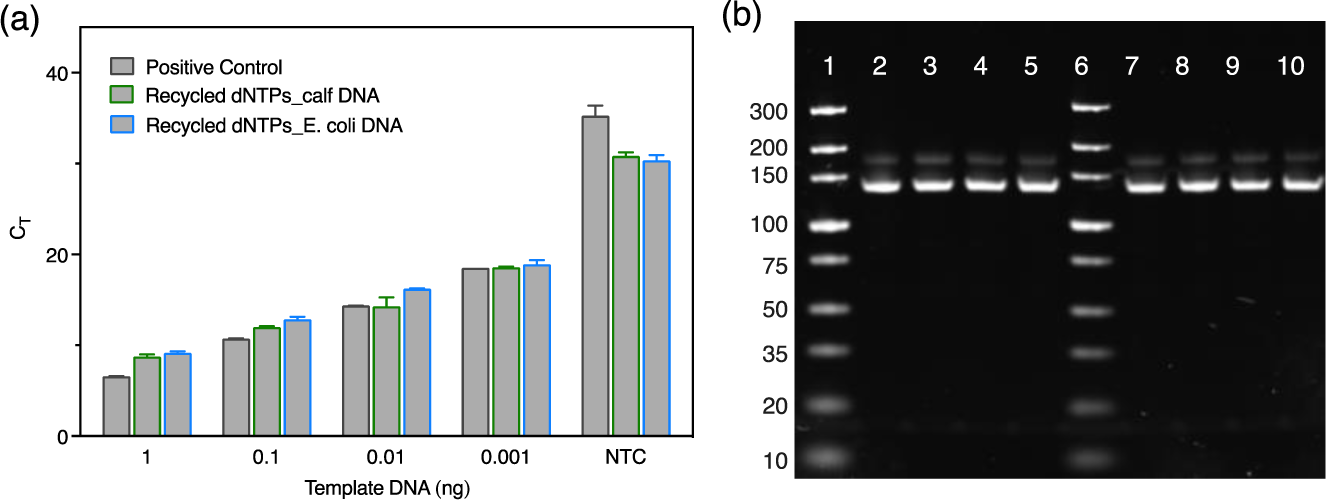
(a) C_T_ values of self-made qPCR kits from recycled dNTPs_calf DNA and dNTPs_E. coli DNA, with commercially available SybrGreen qPCR kit used as positive control. Plots of all qPCR amplification could be found in supporting information. (b) Page-gel of qPCR amplification products (133 base pair), lane 1 and lane 6, ladders; lane 2-5 and lane 7-10, qPCR amplified DNA products. Self-made qPCR kit was prepared from recycled dNTPs_calf DNA (lane 2-5), and from dNTPs_E.coli DNA (lane 7-10). The content of DNA template was 1, 0.1, 0.01, 0.001 ng, respectively.

### Conclusion and outlook

In this work, we have established an efficient approach to recycle DNA. We have shown that DNA can be de-polymerized to generate dNMPs, that in turn can be phosphorylated to generate dNTPs. The so obtained dNTPs can be re-polymerized by using PCR to achieve a new DNA sequence, hence information, that is completely different from the parent one. The obtained dNTPs can also be used as substrate for qPCR with very good DNA detection performance. In addition, this work provides a new top-down method to obtain monomeric nucleotides from Nature available DNA. Since the conventional *de novo* chemical synthesis of nucleotides still has several disadvantages such as multi-steps of synthesis and purification, usage of toxic chemicals and solvents, ^[7,26,27]^ this top-down, one-pot, two-steps of enzymatical hydrolysis and phosphorylation of Nature available DNA could become a totally new path to obtain monomer nucleotides materials with respect to costs, efficiency, reaction condition, and sustainability. Since all the used materials of DNA and enzyme mixtures are abundant in Nature, there is possibility to scale up this process with relatively low cost. Also, during current pandemic situation large-scale of PCR waste is accumulating from the massive covid test, which can be circularly recycled with the same approach. As DNA technology has become a very well-established technique for sequencing, bioengineering, and molecular diagnoses,^[39,40]^ our DNA recycling methodology can be easily adopted to recycle modified nucleotides (e.g., fluorescence labeled dNTPs for sequencing). We believe the established approach has paved the road for the circulation of DNA materials, which can further boost the materials recycling for the development of circular economy, as well as the establishment of related enzymatic reaction systems.

### Supporting Information

Supporting Information is available from the Wiley Online Library or from the author.

## Acknowledgements

This research has been supported by the SNF Spark project CRSK-2_190167, and by the ERC Advanced Grant (884114-NaCRe). We thank the Core Gene Facility (EPFL) for providing the instruments and training of qPCR, and LDPC group (EPFL) for sharing the HPLC. W. Liu thanks Mr. Shiyu Cheng (EPFL_ LBNC group) and for the fruitful discussion, and Ms. Yueyun Zhang (EPFL_UPBRI group) for the help of qPCR experiment.

## Conflicts of interest

This work is undergoing of patent application.

Received: ((will be filled in by the editorial staff))

Revised: ((will be filled in by the editorial staff))

Published online: ((will be filled in by the editorial staff))

## Table of content

Here we show that it is possible to recycle DNA by first hydrolyzing into its monophosphate bases, that in turn are converted into their triphosphate form in a simple one step enzymatic reaction. The triphosphate bases are then used in polymerase chain reaction (PCR) to produce a whole new form of DNA.

ToC figure

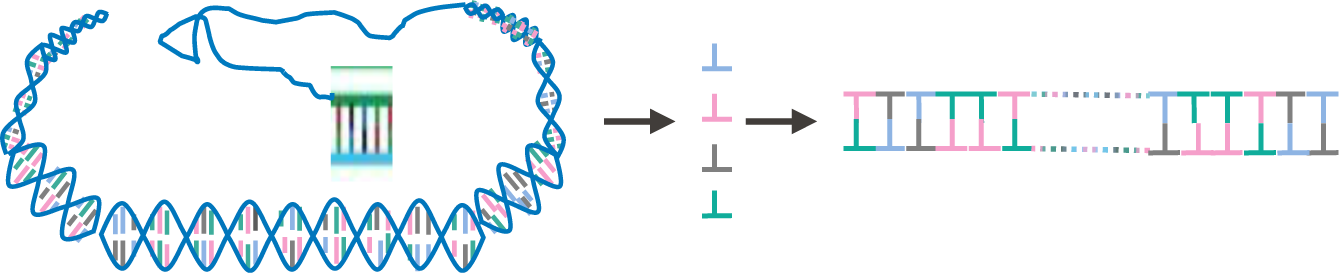

## Supporting Information

### Materials

*Chemicals:* 2’-Deoxyguanosine 5’-monophosphate sodium salt hydrate (dGMP), Thymidine 5′-monophosphate disodium salt hydrate (dTMP), 2′-Deoxycytidine 5′-monophosphate sodium salt (dCMP), 2′-Deoxyadenosine 5′-monophosphate (dAMP), Acetylephosphate Lituium Potassium salt (AceP), Sodium chloride (DNase, RNase, and protease free), Sodium hydroxide solution (5.0 M), Acetic acid, Tetrabutylammonium dihydrogenphosphate solution (1.0 M in water), 2’-Deoxycytidine 5’-diphosphate sodium salt (dCDP), 2’-Deoxyadenosine 5’-diphosiphate sodium salt (dADP), 2’-Deoxyguanosine 5’-diphosphate sodium salt (dGDP), Deoxynucleotide mix reagent (dNTPs, 10 mM for each), Adenosine 5′-diphosphate sodium salt (ADP), Adenosine 5’-triphospate disodium (ATP, 100 mM) were purchased from Sigma-Aldrich. 2’-Deoxythymidine-5’-diphosphate trisodium salt (dTDP) was purchased from abcr. *DNA*: Calf thymus DNA is purchased from Sigma-Aldrich. Cas number 73049-39-5, Catalog number D8661. As described from the vendor, the Calf thymus DNA has been fragmented by sonication. E. coli genomic DNA is purchased from thermo scientific, Cas number 9007-49-2, Catalog number J14380.MA. As described from the vendor, the E. coli genomic DNA is purified from E. coli type B cells, ATCC 11303 strain. It has a single chromosome, and the genome is 4,600,000 bp long. This genomic DNA is fragmented to some degree during purification, yet it is characterized as a high molecular weight DNA. *Enzyme*: Exonuclease III (200 U/µL), Exonuclease I (20 U/µL), DreamTaq DNA Polymerase (5 U/µL) were purchased from Thermo Fisher Scientific. T4 dNMP (deoxy-Nucleotide Monophosphate) Kinase was purchased from Jena Bioscience. E. coli S30 Extract System for Circular DNA (with luciferase plasmid DNA template) was purchased from Promega Corporation. *Chemicals for gel electrophoresis*: 400 µL-SYBR Safe DNA Gel Stain, DNA Loading Dye and SDS Solution (6X), TAE Buffer (Tris-acetate-EDTA) (50x), TrackIt 100 bp DNA Ladder, PowerTrack™ SYBR Green Master Mix were purchased from Thermo Fisher Scientific. Agarose was purchased from Bio-Rad Laboratories. Amicon Ultra-0.5 mL Centrifugal Filters (3 K cutoff) was purchased from Merck. Nuclease-Free Water (10 × 50 ml) was purchased from QIAGEN. *Self-made phosphorylation buffer:* 55mM HEPES, Magnesium acetate 15 mM, pH 7.5. HEPES buffer (1 M) was purchased from Thermo Fisher Scientific. NuPAGE 4-12% Bis-Tris Protein Gels, NuPAGE™ MES SDS Running Buffer (20X) were purchased from Thermo Fisher Scientific. Details of materials, experiments, and instrument information about GFP plasmid DNA amplification and GFP expression in experiment section.

### Instruments

Eppendorf Thermomixer (RTM F1.5, 220 - 240 V/50 - 60 Hz) was purchased from Eppendorf. DNA ultrasonication was performed by Vibra-Cell™ 75286 ultrasonic Liquid Processors. Horizontal gel electrophoresis system was purchased from Bio-Rad. Gel images was taken from GelDoc Go, Bio-rad. PCR was performed by Proflex 3×32-well PCR thermal cycler system (ThermoFisher Scientific). Quantitative PCR was performed by QuantStudio 7 qPCR instrument (Applied Biosystems). HPLC was performed by Infinite 1260 HPLC with C 18 column, Agilent. Details of LC-MS system in the experiment section.

### Experiment

#### 1. DNA hydrolysis

##### Calf thymus DNA hydrolysis and HPLC characterization

Calf thymus DNA (sheared, 0.966 mg/mL, 75 µL) was mixed with 3 µL Exonuclease III (600 Unit), 3 µL Exonuclease I (60 Unit), 9 µL nuclease free water, and 10 µL 10 x Exonuclease III buffer. The final reaction volume is 100 µL with 1 x Exonuclease III buffer (0.66 mM MgCl_2_, 66 mM Tris-HCl, pH 8.0 at 30 °C). The Calf DNA hydrolysis mixture (referred as **dNMPs_calf DNA-1**) was incubated in thermomixer at 37°C, 350 RPM overnight. Following the DNA hydrolysis mixture was incubated in 80°C for 20 mins to inactivate the Nuclease. The dNMPs recycled from Calf DNA was characterized by HPLC. *As the retention peaks of dGMP and dTMP were not very well separated (Figure S3), the concentration of dNMPs was quantified by LC-MS*.

##### Calf thymus DNA hydrolysis and LC-MS characterization

Calf thymus DNA (sheared, 2.5 mg/mL, 150 µL) was mixed with 15 µL Exonuclease III (3000 Unit), 15 µL Exonuclease I (300 Unit), 90 µL nuclease free water, and 30 µL 10 X Exonuclease III buffer. The final reaction volume is 300 µL with 1 X Exonuclease III buffer (0.66 mM MgCl_2_, 66 mM Tris-HCl, pH 8.0 at 30 °C). The Calf DNA hydrolysis mixture was incubated in thermomixer at 37°C, 350 RPM overnight. Following the DNA hydrolysis mixture was incubated in 80°C for 20 mins to inactivate the Nuclease. The dNMPs recycled from Calf DNA was referred as **dNMPs_calf DNA-2** and quantified by LC-MS (Figure S1b). DNA hydrolysis substrate was prepared as dilution buffer for the dNMPs standard solution. Reaction mixture of DNA hydrolysis buffer without calf DNA (with substrate condition equal to the DNA hydrolysis condition) was prepared by mixing 3 µL Exonuclease III, 3 µL Exonuclease I, 48 µL nuclease free water, and 6 µL 10 X Exonuclease III buffer. The reaction mixture was purified by ultrafiltration (Amicon, 3KD), diluted for 5000 times, and later used as dilution buffer for the preparation of dNMPs standard solution. Stock solution of dNMPs standard (1 mM for each) was prepared in milli-Q water. Further the stock solution was stepwisely diluted to 1000, 800, 600, 400, 200, 100, 50 nM by the diluted DNA hydrolysis buffer. The reason to prepare dNMPs standard solution in DNA hydrolysis buffer, is to maintain the sample and references in the same condition, so that the influences from ion suppression for LC-MS quantification results can be avoided.

#### 2. dNMPs Phosphorylation

##### dNMPs-phosphorylation and HPLC quantification

The dNMPs (100 mM, 0.8 µL for each) was mixed with 2 µL E. coli S30 Extract, 0.66 µL T4 dNMP Kinase (66 Unit), 10 µL ATP (10 mM), and 25.6 µL Acetyl phosphate Lithium potassium (AceP, 50 mM), 20 µL phosphorylation buffer, and 138.54 µL nuclease-free water for phosphorylation. The dNMPs_phosphorylation reaction mixture with final volume 200 µL, dNMPs (0.4 mM for each), ATP (0.5 mM), AceP (6.4 mM, 2 equivalent) was incubated in thermomixer at 400 RPM, 37°C for 4 hours. All the hydrolysis and phosphorylation enzymes were removed by ultrafiltration (Amicon, 3 KD cutoff). Further the filtrated reaction mixture referred as **dNTPs_dNMPs** was diluted for 50 times and injected to HPLC (50 µL) to evaluate the phosphorylation efficiency.

##### dNMPs_calf DNA phosphorylation and HPLC quantification

The hydrolyzed Calf DNA (**dNMPs_calf DNA-1**, 66.7 µL) was mixed with 1 µL E. coli S30 Extract, 1 µL T4 NMP Kinase (3 times dilution, 33 Unit), 5 µL ATP (10 mM), and 12.8 µL Acetyl phosphate Lithium potassium (AceP, 50 mM), 10 µL phosphorylation buffer, 3.5 µL nuclease-free water for phosphorylation. The dNMPs_calf DNA phosphorylation reaction mixture with final volume 100 µL, estimated dNMPs (in average 0.4 mM for each), ATP (0.5 mM), AceP (6.4 mM, 2 equivalent) was incubated in thermomixer at 400 RPM, 37°C for 4 hours. Afterwards all hydrolysis enzymes and phosphorylation enzymes, and non-hydrolyzed DNA was removed by ultrafiltration (Amicon, 3 KD cutoff, 5000 RPM for 30 min in 4°C). Further the filtrated reaction mixture (referred as **dNTPs_calf DNA-1)** was diluted for 50 times and injected to HPLC (50 µL) for quantification of each dNTP. The recycled nucleotide mixture **dNTPs_calf DNA-1** was directly used for PCR to amplify GFP DNA plasmid.

#### 3. LC-MS

The LC-MS characterization of recycled dNMPs experiments were carried out using a shorter version of the protocol published by Zhang et at.^[1]^ Analysis were conducted on a Xevo G2-S QTOF mass spectrometer coupled to the Acquity UPLC Class Binary Solvent manager and BTN sample manager (Waters, Corporation, Milford, MA). The injection volume was 5 μL. Mass spectrometer detection was operated in negative ionization using the ZSpray™ dual-orthogonal multimode ESI/APCI/ESCi® source. The TOF mass spectra were acquired in the resolution mode over the range of m/z 100-500 at an acquisition rate of 0.1 sec/spectra. The instrument was calibrated using a solution of sodium format (0.01 mg/L in isopropanol/H2O 90:10). A mass accuracy better than 5 ppm was achieved using a Leucine Enkephalin solution as lock-mass (200 pg/mL in ACN/H2O (50:50)) infused continuously using the LockSpray source. Source settings were as follows: cone, 25V; capillary, 3 kV, source temperature, 140°C; desolation temperature, 400°C, cone gas, 70 L/h, desolation gas, 500 L/h. Data were processed using MassLynx™ 4.1 software and QuanLynx application for quantification. The separation was achieved using an ACQUITY Premier HSS 1.8 µm vanguard FIT column, 2.1 mm x 100 mm (Waters) heated at 40°C. Mobile phase consisted of 0.1% formic acid in water as eluent A and 0.1% formic acid in acetonitrile/water (6:4) as eluent B. The separation was carried out at mL/min over a 15 min total run time using the following program: from 0 to 5 min, 100-95% A; 5-10 min, 95% A; 10-10.1 min, 95-100 % A; 10.1-15 min, 100% B to re-equilibrate the system in initial conditions.

Standard stock solutions of dNMPs mixture were prepared at a concentration of 1 mM in Milli-Q water. Stock solutions were further diluted in DNA hydrolysis buffer (5000 times diluted) and calibration curves achieved by a serial dilution in the 50–1000 nM concentration range (Figure S1a). The DNA hydrolysis product **dNMPs_calf DNA-2** were diluted in Milli-Q water for 5000 times before LC-MS analysis in order to fit into the calibration curves. Extracted ions chromatograms (XIC) were based on a retention time (RT) window of ± 0.25 min with a mass-extraction-window (MEW) of ± 25 ppm centered on m/z of each nucleotide. The average peak area of three replicate injections at each concentration was used for each data point. Calibration curves were fitted with a polynomial order 2 equation, with R2 > 0.98 for all nucleotides.

**Figure S1.**
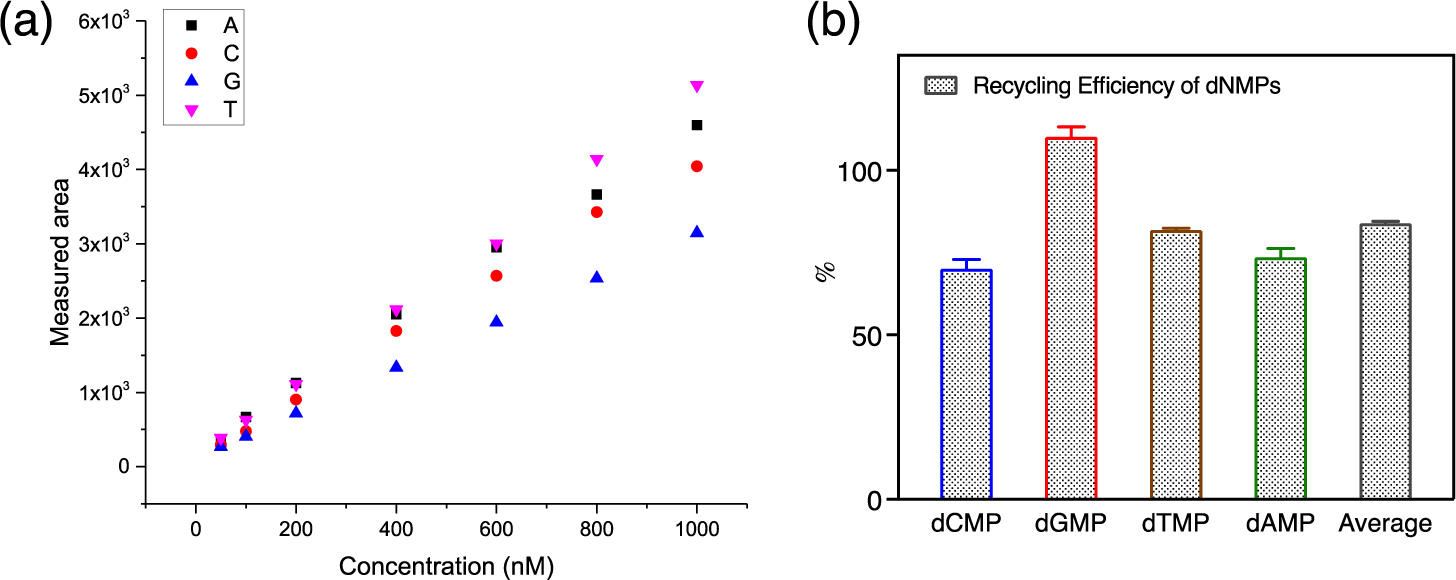
**(a)** Calibration curve of dNMPs from LC-MS, concentration (50, 100, 200, 400, 600, 800, 1000 nM). (b) Recycling rate of dNMPs from calf DNA.

#### 4. HPLC quantification

The concentration of dNTPs was quantified by HPLC with C 18 column. Mobile phase Buffer A: 5 mM t-butyl ammonium phosphate, 10 mM KH_2_PO_4_, and 0.25% methanol adjusted to pH 6.9. Buffer B: 5 mM t-butyl ammonium phosphate, 50 mM KH_2_PO_4_, and 30% methanol (pH 7.0). From 0 to15 mins gradients changed from 40%/60% to 20%/80% of buffer A/B and run under the same gradient condition to 20 min, and changed back to the starting condition of 40%/60%, flow rate 0.5 ml/min.

Mixture of dNMPs (8 µM for each, 50 µL) was injected to HPLC. The retention time of nucleotide monophosphate are as following: dCMP_3.4 mins, dGMP_4.2 mins, dTMP_4.6 mins, dAMP_6.7 mins. Mixture of dNTPs and ATP (2.5, 5, 10, 20, 40 µM) standard solution was injected to HPLC to generate the calibration curves for quantification (Figure S3). Calibration curve of dNTPs as well as ATP was generated by integrated area. The retention time of nucleotide triphosphate (dNTPs and ATP) are as following: dCTP_9.5 mins, dGTP_12.4 mins, dTTP_14.3 mins, ATP_15.7 mins, dATP_18.2 mins (Figure S2). Phosphorylation products of **dNTPs_dNMPs** and **dNTPs_calf DNA-1** were injected to HPLC for quantification.

**Figure S2.**
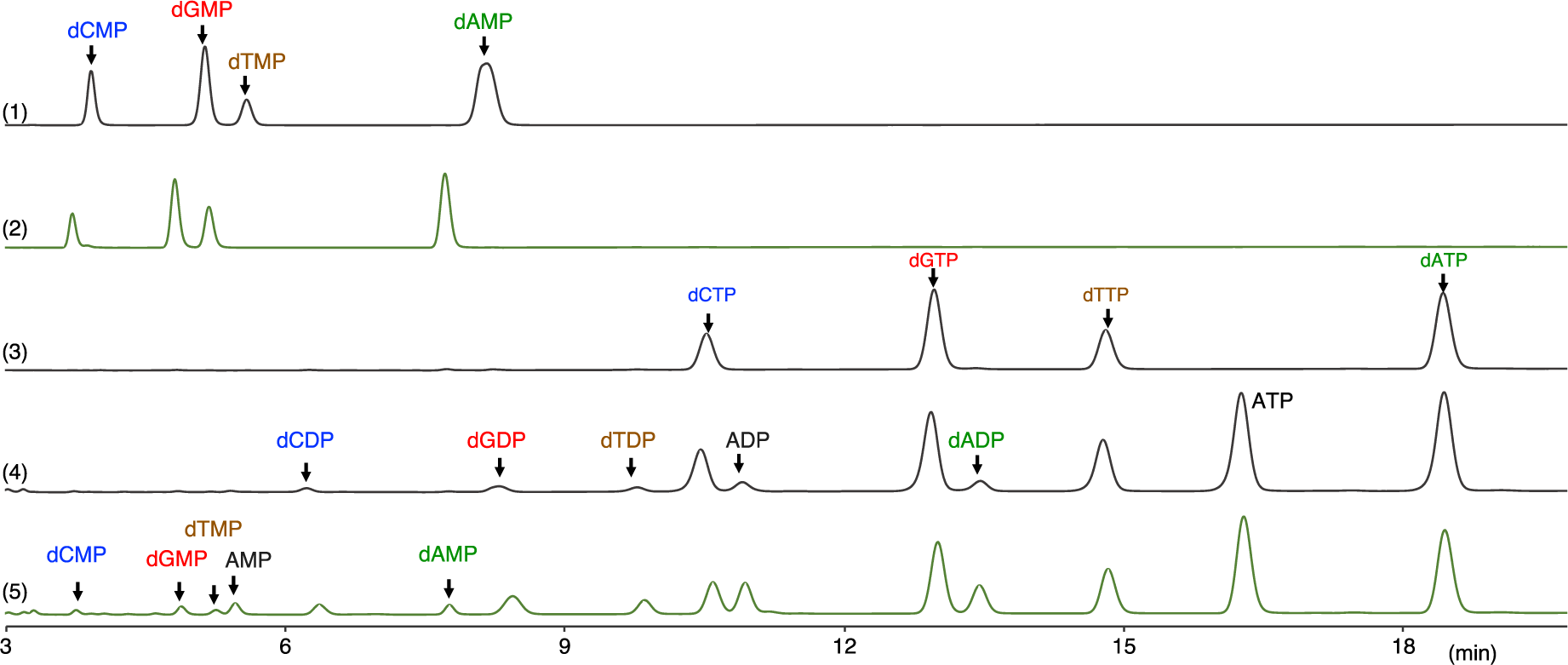
HPLC retention time of (1) dNMPs standards, (2) dNMPs_calf DNA, (3) dNTPs standards, (4) dNTPs_dNMPs phosphorylation, and (5) dNTPs_calf DNA.

**Figure S3.**
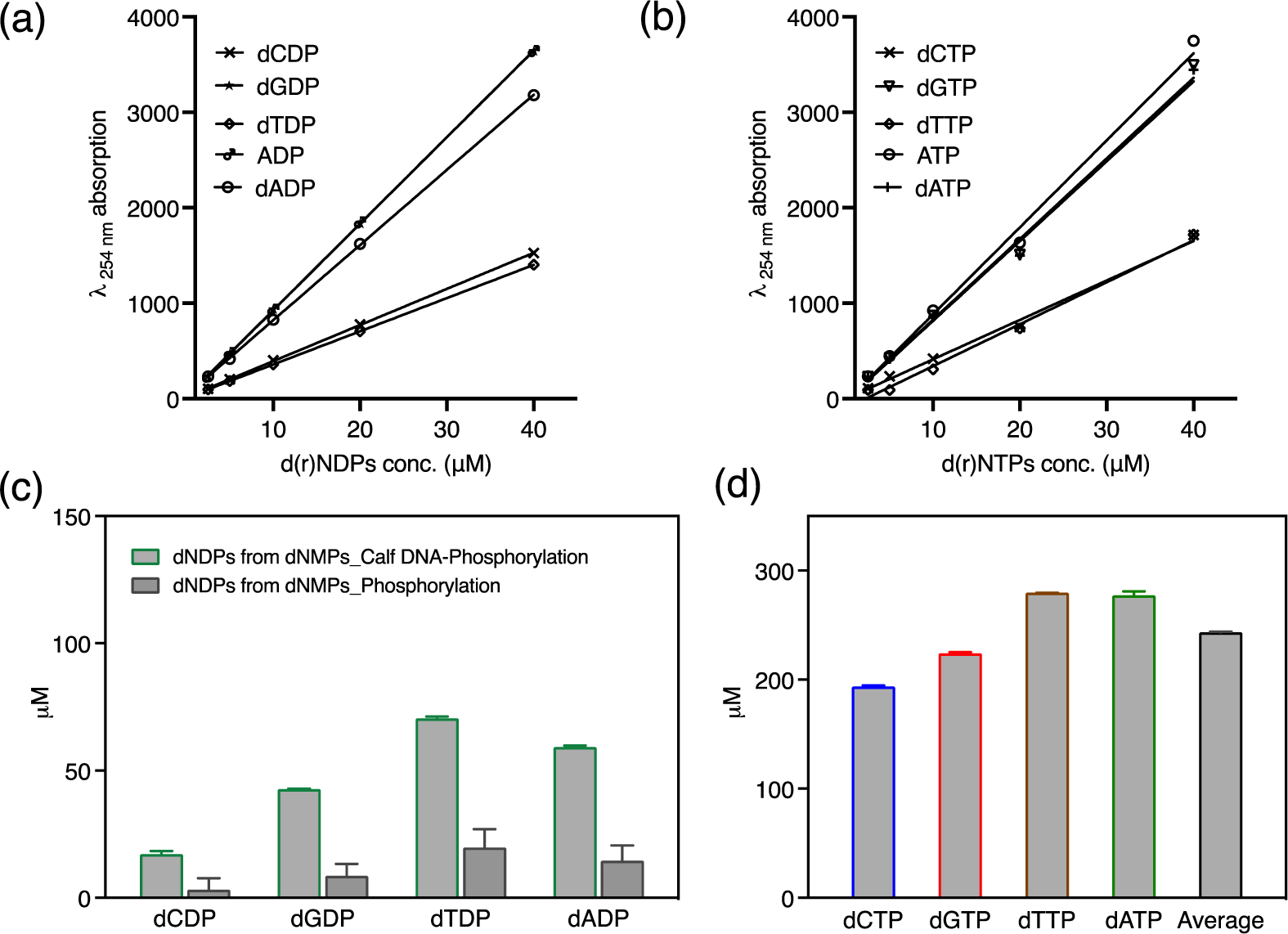
(a) Calibration curve of d(r)NDPs (concentration 2.5, 5, 10, 20, 40 µM). (b) Calibration curve of dNTPs and ATP standard solution (concentration 2.5, 5, 10, 20, 40 µM). (c)Concentration of dNDPs residues from the phosphorylation products of dNTPs_calf DNA (dCDP 17.13 ± 1.26 µM, dGDP 42.68 ± 0.19 µM, dTDP 70.46 ± 0.74 µM, dADP 59.20 ± 0.63 µM), and from dNMPs phosphorylation reaction (dCDP 3.17 ± 4.49 µM, dGDP 8.60 ± 4.68 µM, dTDP 19.72 ± 7.25 µM, dADP 14.53 ± 6.01 µM). (d) Concentration of dNTPs recycled from calf DNA (dCTP 193.7 ± 0.8 µM, dGTP 224.1 ± 0.9 µM, dTTP 279.5 ± 0.1 µM, dATP 277.3 ± 3.7 µM, in average 243.7 ± 0.9 µM).

#### 5. Gel electrophoreses

Samples of calf DNA (before and after hydrolysis) were mixed with DNA loading dye and loaded to an agarose gel (2%). The loading amount was adjusted to equal amount of calf DNA before and after hydrolysis. The agarose gel was run in 1 x TAE buffer at 120 V for 40 min. Afterwards, the gel was stained by 1x Sybr safe solution for 40 min under slow shaking. Following the gel image was taken by GelDoc Go under Sybr safe channel for 1 s exposure time (Figure 1b). lane 1, TrackIt 100 bp ladder, 2 µL; lane 2, calf DNA, 0.966 mg/mL, 2.1 µL; lane 3, hydrolyzed calf DNA, 2.73 µL (1.3 x dilution by hydrolysis).

#### 6. Recycled dNTPs for GFP DNA amplification and GFP expression Materials

*PCR reagents*. gBlock encoding GFP, and primers (fwd and rev) were purchased from IDT Integrated DNA Technologies. 5x Phusion HF Buffer, dNTP Mix (10 mM), Phusion High-Fidelity DNA Polymerase (2 U μl^−1^), and DMSO were purchased from Thermo Fisher Scientific; nuclease-free water was supplied by Sigma-Aldrich. 5x GelPilot DNA Loading Dye, and QIAquick PCR Purification Kit were purchased from Qiagen; GeneRuler 1 kb DNA Ladder (ready-to-use), and SYBR Safe DNA Gel Stain from Thermo Fisher Scientific. UltraPure Agarose was supplied by Invitrogen. 50x TAE buffer was purchased from Jena Bioscience. *Cell-Free expression*. Magnesium acetate, Potassium glutamate, DL-Dithiothreitol (DTT), Creatine phosphate, Folinic acid, Spermidine, HEPES buffer, Protector RNase Inhibitor, and the 20 proteinogenic AAs were purchased from Sigma-Aldrich. ATP, GTP, CTP, and UTP were supplied by Thermo Fisher Scientific. tRNAs were purchased from Roche. PUREfrex™ Solution II (enzymes), and PUREfrex™ Solution III (ribosomes) were supplied by Kaneka Eurogentec SA. *Tools*. Protein LoBind Tubes were purchased from Eppendorf. Nunc™ 384-well optical bottom plates were supplied by Thermo Fisher Scientific. SealPlate sealing film was purchased from Sigma-Aldrich.

##### Recycled dNTPs for GFP plasmid DNA amplification

*Positive control PCR batch (20 μl)*. The reaction was assembled by mixing 1 μl DNA linear gBlock template (1 ng μl^−1^), 0.2 μl fwd. primer (50 μM), 0.2 μl rev. primer (50 μM), 4 μl 5x Phusion HF Buffer, 0.3 μl dNTP Mix (10 mM), 1 μl DMSO, 0.15 μl Phusion High-Fidelity DNA Polymerase (2 U μl^−1^), and 13.15 μl nuclease-free water in a small PCR vial. *Sample PCR batch (20 μl)*. The reaction was assembled by mixing 1 μl DNA linear gBlock template (1 ng μl^−1^), 0.2 μl fwd. primer (50 μM), 0.2 μl rev. primer (50 μM), 4 μl 5x Phusion HF Buffer, 12.5 μl dNTP Mix (0.24 mM), 1 μl DMSO, 0.15 μl Phusion High-Fidelity DNA Polymerase (2 U μl^−1^), and 0.95 μl nuclease-free water in a small PCR vial. *PCR thermal cycle (20 μl batch)*. Initialization was run at 98° C for 2 min, denaturation at 98° C for 20 s, annealing at 47° C for 30 s, and extension at 72° C for 45 s. Denaturation, annealing, and extension were repeated 35x. The reaction temperature was kept at 72° C for additional 7 min and decreased to 4° C for storage. The whole thermal cycle was run into Thermo Fisher Scientific ProFlex™ PCR System. *Casting of the gel*. The size of the amplified template was checked by running an agarose gel, prior to purification of the template from the PCR batch. 1% Agarose gel was cast by mixing g of Agarose into 40 ml of 1x TAE buffer; the suspension was heated in the microwave at 800 W for 90 s approximately and added with 4 μl of SYBR Safe DNA Gel Stain. *Samples preparation*. 1 μl of PCR reaction was diluted adding 3 μl of nuclease-free water, and 1 μl of 5x GelPilot DNA Loading Dye; 5 μl of GeneRuler 1 kb DNA Ladder were used as reference. *Running conditions*. The gel was run at 60 V for 5 min followed by 120 V for 30 min in the Thermo Scientific EasyCast gel system. *Imaging*. The gel was imaged by using Thermo Fisher Scientific Benchtop 3UV transilluminator equipped with Kodak gel logic 100 imaging system, λ = 302 nm, 4s exposure. *Purification*. The PCR product was purified by combining 4 PCR batches, doubling the final volume by adding nuclease-free water, and following the QIAquick PCR Purification Kit protocol. DNA was eluted by using 15 μl of elution buffer per spin column. The final DNA concentration was measured using Witec NanoDrop 1000 spectrophotometer.

##### Cell-free protein TX-TL

*Energy solution preparation*. The following solutions were prepared. SolutionA(-Salts – tRNAs - AAs) (2 ml): Creatine phosphate (147.06 mM), Folinic acid (0.15 mM), Spermidine (14.71 mM), DTT (7.4 mM), ATP (14.71 mM), GTP (14.71 mM), CTP (7.4 mM), UTP (7.4 mM), and HEPES (pH 7.6, 367.65 mM). Salts solution (2 ml): Magnesium acetate (184.38 mM), and Potassium glutamate (1.563 M). tRNAs solution (200 μl): tRNAs (560 A_260_ mL^−1^). tRNAs were quantified by using UV absorption A_260_ in Witec NanoDrop 1000 spectrophotometer. The three solutions were combined in a 25 μl reaction, by mixing 3.4/1.6/2.5 v/v/v solutionA(-Salts - tRNAs - AAs):salts solution:tRNAs solution, in order to get the desired concentrations, adapted from Ueda and coworkers:^[2]^ Creatine phosphate (20 mM), Folinic acid (0.02 mM), Spermidine (2 mM), DTT (1 mM), ATP (2 mM), GTP (2 mM), CTP (1 mM), UTP (1 mM), HEPES (pH 7.6, 50 mM), Magnesium acetate (11.8 mM), Potassium glutamate (100 mM), and tRNAs (56 A_260_ ml^−1^). *Cell-Free TX-TL reactions assembly (25 μl)*. 3.4 μl of solutionA(-Salts - tRNAs - AAs), 1.6 μl of salts solution, 2.5 μl of tRNAs solution, 1.25 μl PUREfrex™ Solution II (enzymes), 1.25 μl PUREfrex™ Solution III (ribosomes), 0.5 μl RNAse inhibitor, 2.5 μl of AAs, and 75 ng DNA (samples, and positive controls) were mixed in ice. Nuclease-free water was added to bring the reaction volume to 25 μl. In the negative controls the DNA was replaced with nuclease-free water. These volumes keep each reagent at the desired concentration in the TX-TL reaction. *Cell-Free TX-TL reaction*. The reactions were gently mixed, transferred into a 384-well plate, sealed to avoid evaporation, spin down at 3000 rcf, 25° C in Eppendorf 5810R, and incubated at 37° C for 8 h in Thermo Fisher Scientific BioTek Synergy Mx plate reader. The plate reader parameters were the following: detection method = fluorescence, λ_exc_ = 488 nm, λ_em_ = 507 nm, 1 min interval read, sensitivity = 70 %, bottom optic position, fast continuous shaking. *Data processing*. The TX-TL reactions were all run in duplicates. The expression curves represent the statistical mean of the results at any acquisition time; the shadow represents the standard deviation of the same data.

##### DNA sequences gBlock (GFP)

(5’)gcaccatcagccagaaaaccgaaccagccagaaaacgacctttctgtggatcttaaggctagagtactaatacgactcactatagggagaccacaacggtttccctctagaaataattttgtttaacttaagaaggaggaaaaaaaaatggtctctaaaggtgaagaattattcactggtgttgtcccaattttggttgaattagatggtgatgttaatggtcacaaattttctgtctccggtgaaggtgaaggtgatgctacttacggtaaattgaccttaaaatttatttgtactactggtaaattgccagttccatggccaaccttagtcactactttaacttatggtgttcaatgtttttctagatacccagatcatatgaaacaacatgactttttcaagtctgccatgccagaaggttatgttcaagaaagaactatttttttcaaagatgacggtaactacaagaccagagctgaagtcaagtttgaaggtgataccttagttaatagaatcgaattaaaaggtattgattttaaagaagatggtaacattttaggtcacaaattggaatacaactataactctcacaatgtttacatcatggctgacaaacaaaagaatggtatcaaagttaacttcaaaattagacacaacattgaagatggttctgttcaattagctgaccattatcaacaaaatactccaattggtgatggtccagtcttgttaccagacaaccattacttatccactcaatctgccttatccaaagatccaaacgaaaagagagaccacatggtcttgttagaatttgttactgctgctggtattaccttaggtatggatgaattgtacaaacaccaccatcatcaccactaataacgactcaggctgctacctagcataaccccttggggcctctaaacgggtcttgaggggttttttggcaggaaagaacatgtgagcaaaagg(3’)

Forward primer: 5’-gatcttaaggctagagtac-3’

Reverse primer: 5’-caaaaaacccctcaagac-3’

##### Protein Sequentes (GFP)

MVSKGEELFTGVVPILVELDGDVNGHKFSVSGEGEGDATYGKLTLKFICTTGKLPVPWPTLVTTLTYGVQCFSRYPDHMKQHDFFKSAMPEGYVQERTIFFKDDGNYKTRAEVKFEGDTLVNRIELKGIDFKEDGNILGHKLEYNYNSHNVYIMADKQKNGIKVNFKIRHNIEDGSVQLADHYQQNTPIGDGPVLLPDNHYLSTQSALSKDPNEKRDHMVLLEFVTAAGITLGMDELYKHHHHHH&

#### 7. Recycle dNTPs for quantitative PCR (qPCR)

Based on the quantification of hydrolysis yield (83.9%) and phosphorylation yield (69.0%) of DNA recycling process, the DNA recycling condition was adjusted accordingly to obtain recycled dNTPs with estimated final average concentration 0.4 mM for each for qPCR.

##### dNTPs recycling from calf DNA

Calf DNA (2.5 mg/mL, 150 µL) was mixed with 15 µL Exonuclease III (3000 Unit), 15 µL Exonuclease I (300 Unit), 90 µL nuclease free water, and 30 µL 10 X Exonuclease III buffer. The final reaction volume is 300 µL with 1 X Exonuclease III buffer (0.66 mM MgCl_2_, 66 mM Tris-HCl, pH 8.0 at 30 °C). The DNA hydrolysis mixture (dNMPs_calf DNA-3) was incubated in thermomixer at 37°C, 350 RPM overnight, and following incubation in 80°C for 20 min to inactivate the Nuclease. The DNA hydrolysis mixture referred as **dNMPs_calf DNA-3** (134 µL) was mixed with 2 µL E. coli S30 Extract, 0.66 µL T4 dNMP Kinase (66 Unit), 10 µL ATP (10 mM), and 25.6 µL Acetyl phosphate Lithium potassium (AceP, 50 mM), 20 µL phosphorylation buffer, and 7.74 µL nuclease-free water for phosphorylation. The phosphorylation reaction mixture with final volume 200 µL, with ATP (0.5 mM), AceP (6.4 mM, 2 equivalent), and estimated final concentration of dNTPs (0.4 mM in average), was incubated in thermomixer at 400 RPM, 37°C for 4 hours. Afterwards all the hydrolysis and phosphorylation enzymes were removed by ultrafiltration (Amicon, 3 KD cutoff). Further the filtrated reaction mixture referred as **dNTPs_calf DNA-3** was directly applied for qPCR. The Mg^2+^ concentration originated from hydrolysis and phosphorylation buffer as well as storage condition of enzymes was calculated with final concentration of 1.55 mM.

##### dNTPs recycling from E. coli DNA

Non-sheared E. coli DNA was purchased from thermo scientific. The E. coli DNA (5 mg/mL, 2 mL in a 5 mL glass vial) was firstly sheared by ultrasonication for 30 mins (10 seconds of 90% amplitude + 5 seconds pause) in ice bath. Following the sheared E. coli DNA (2.5 mg/mL, 150 µL) was mixed with 15 µL Exonuclease III (3000 Unit), 15 µL Exonuclease I (300 Unit), 90 µL nuclease free water, and 30 µL 10 X Exonuclease III buffer. The final reaction volume is 300 µL with 1 X Exonuclease III buffer (0.66 mM MgCl_2_, 66 mM Tris-HCl, pH 8.0 at 30 °C). The DNA hydrolysis mixture (dNMPs_E. coli DNA-1) was incubated in thermomixer at 37°C, 350 RPM overnight, and following incubation in 80°C for 20 min to inactivate the Nuclease. Further the DNA hydrolysis mixture referred as **dNMPs_E. coli DNA** (134 µL) was mixed with 2 µL E. coli S30 Extract, 0.66 µL T4 dNMP Kinase (66 Unit), 10 µL ATP (10 mM), and 25.6 µL Acetyl phosphate Lithium potassium (AceP, 50 mM), 20 µL phosphorylation buffer, and 7.74 µL nuclease-free water for phosphorylation. The reaction mixture with final volume 200 µL, ATP (0.5 mM), AceP (6.4 mM, 2 equivalent), and estimated final concentration of dNTPs (0.4 mM in average), was incubated in thermomixer at 400 RPM, 37°C for 4 hours. Afterwards all the hydrolysis and phosphorylation enzymes were removed by ultrafiltration (Amicon, 3 KD cutoff). Further the filtrated reaction mixture referred as **dNTPs_E. coli DNA** was directly applied for qPCR. The Mg^2+^ concentration originated from hydrolysis and phosphorylation buffer as well as storage condition of enzymes was calculated with final concentration of 1.55 mM.

##### qPCR materials

Plasmid luciferase DNA (4864 bp) was from the E. coli S30 extraction kit with luciferase sequence (pBEST luc, 1684 bp). Primers were designed by IDT PrimerQuest™ Tool with forward primer starts from 673 of luciferase sequence, reverse primer start from 805 of luciferase sequence, and length of amplicon 133 base pair. Primers were ordered from Biomers Gmbh.

qPCR reaction mixture of recycled dNTPs was prepared as following: *qPCR_Calf DNA:* dNTPs_calf DNA-3 (0.4 mM for each, 100 µL), DreamTaq polymerase (1 µL, 2.5 U), 10 X DreamTaq buffer (10 µL), Sybr Dye 1000 x (20 µL), Nuclease water (9 µL) in total 140 µL. *qPCR_E. coli DNA:* dNTPs_E. coli DNA (0.4 mM for each, 100 µL), DreamTaq polymerase (1 µL, 5 U), 10x DreamTaq buffer (10 µL), Sybr Dye 1000x (20 µL), Nuclease water (9 µL) in total 140 µL.

Forward primer: 5’-cgc atg cca gag atc cta tt -3’

Reverse primer: 5’-aga cga ctc gaa atc cac ata tc -3’

qPCR kits (duplicate) were prepared by recycled dNTPs as following:

| Component | Stock conc. | Volume for 1 reaction | Final conc. |
| --- | --- | --- | --- |
| Luc template DNA | 1 ng/ μL,<br>0.1 ng/ μL,<br>0.01 ng/ μL,<br>0.001 ng/ μL,<br>0 ng/ μL, NTC | 1 μL,<br>1 μL,<br>1 μL,<br>1 μL,<br>1 μL | 1 ng,<br>0.1 ng,<br>0.01 ng,<br>0.001 ng,<br>0 ng for NTC |
| qPCR_calf DNA<br>or qPCR_E. coli DNA | 0.285 mM for each | 7 μL | 0.2 mM for each |
| Primer (for. and rev.) | 2.5 μM for each | 1.6 μL | 400 nM |
| Nuclease-free water | | 0.4 μL | |
| Total | 10 μL | 10 μL | 10 μL |

Positive control, duplicate

| Component | Stock conc. | Volume for 1 reaction | Final conc. |
| --- | --- | --- | --- |
| Luc template DNA | 1 ng/ μL,<br>0.1 ng/ μL,<br>0.01 ng/ μL,<br>0.001 ng/ μL,<br>0 ng/ μL, NTC | 1 μL,<br>1 μL,<br>1 μL,<br>1 μL,<br>1 μL | 1 ng,<br>0.1 ng,<br>0.01 ng,<br>0.001 ng,<br>0 ng for NTC |
| qPCR master mix | 0.4 mM for each | 5 μL | 0.2 mM for each |
| Primer (for. and rev.) | 2.5 μM for each | 1.6 μL | 400 nM |
| Nuclease-free water | | 2.4 μL | |
| Total | 10 μL | 10 μL | 10 μL |

##### Thermocycle condition for qPCR

the qPCR amplification was performed by QuantStudio 7 qPCR system using the following thermal cycling conditions: Initial denature (95°C, 2 min) for 1 cycle, amplification (denature at 95°C for 15s, annealing and amplification at 60 °C for 30 s) for 40 cycles.

**Figure S4.**
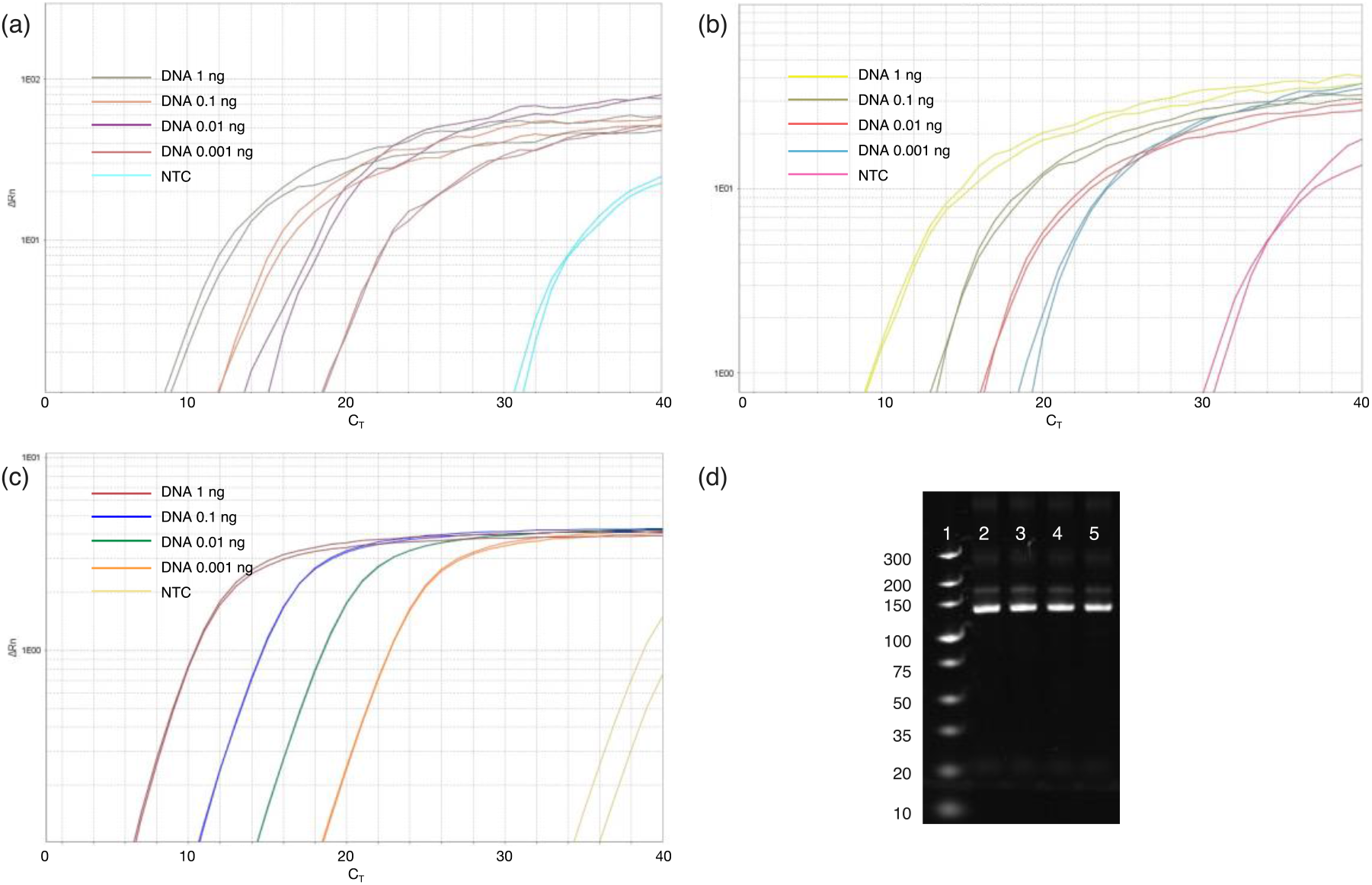
Amplification plots of self-made qPCR kits from dNTPs_calf DNA (a), dNTPs_E. coli DNA (b), and commercial qPCR kit as positive control (c), with DNA template 1, 0.1, 0.01, 0.001 ng and NTC, respectively. (d) Page-gel of qPCR amplification product from positive control with DNA template 1, 0.1, 0.01, 0.001 ng (lane 2-5).

To test if the residue of dNDPs and ATP in the self-made qPCR kit would affect the qPCR performance, qPCR assay was prepared with/without 0.2 mM ATP or 6.25 - 100 µM dNDPs added. There was no obvious change of the C_T_ values with ATP added (Figure S5a and S5b), and only slightly change of the C_T_ values (less than 1 cicle) with higher concentration of dNDPs (Figure S5c and S5d).

qPCR with or without ATP, duplicate

| Component | Stock conc. | Volume for 1 reaction | Final conc. |
| --- | --- | --- | --- |
| Luc template DNA | 1 ng/ μL,<br>0.1 ng/ μL,<br>0.01 ng/ μL,<br>0.001 ng/ μL,<br>0 ng/ μL, NTC | 1 μL,<br>1 μL,<br>1 μL,<br>1 μL,<br>1 μL | 1 ng,<br>0.1 ng,<br>0.01 ng,<br>0.001 ng,<br>0 ng for NTC |
| qPCR master mix | 0.4 mM for each | 5 μL | 0.2 mM for each |
| ATP | 2 mM | 0 or 1 μL | 0 or 0.2 mM for each |
| Primer (for. and rev.) | 2.5 μM for each | 1.6 μL | 400 nM |
| Nuclease-free water | | 2.4 or 1.4 μL | |
| Total | 10 μL | 10 μL | 10 μL |

qPCR with or without dNDPs, duplicate

| Component | Stock conc. | Volume for 1 reaction | Final conc. |
| --- | --- | --- | --- |
| Luc template DNA | 1 ng/ μL,<br>0.1 ng/ μL,<br>0.01 ng/ μL,<br>0.001 ng/ μL,<br>0 ng/ μL, NTC | 1 μL,<br>1 μL,<br>1 μL,<br>1 μL,<br>1 μL | 1 ng,<br>0.1 ng,<br>0.01 ng,<br>0.001 ng,<br>0 ng for NTC |
| qPCR master mix | 0.4 mM for each | 5 μL | 0.2 mM for each |
| dNDPs | 0 μM<br>31.25 μM<br>62.5 μM<br>125 μM<br>250 μM<br>500 μM | 2 μL,<br>2 μL,<br>2 μL,<br>2 μL,<br>2 μL,<br>2 μL | 0<br>6.25 μM for each,<br>12.5 μM for each,<br>25 μM for each,<br>50 μM for each,<br>100 μM for each |
| Primer (for. and rev.) | 2.5 μM for each | 1.6 μL | 400 nM |
| Nuclease-free water | | 0.4 μL | |
| Total | 10 μL | 10 μL | 10 μL |

**Figure S5.**
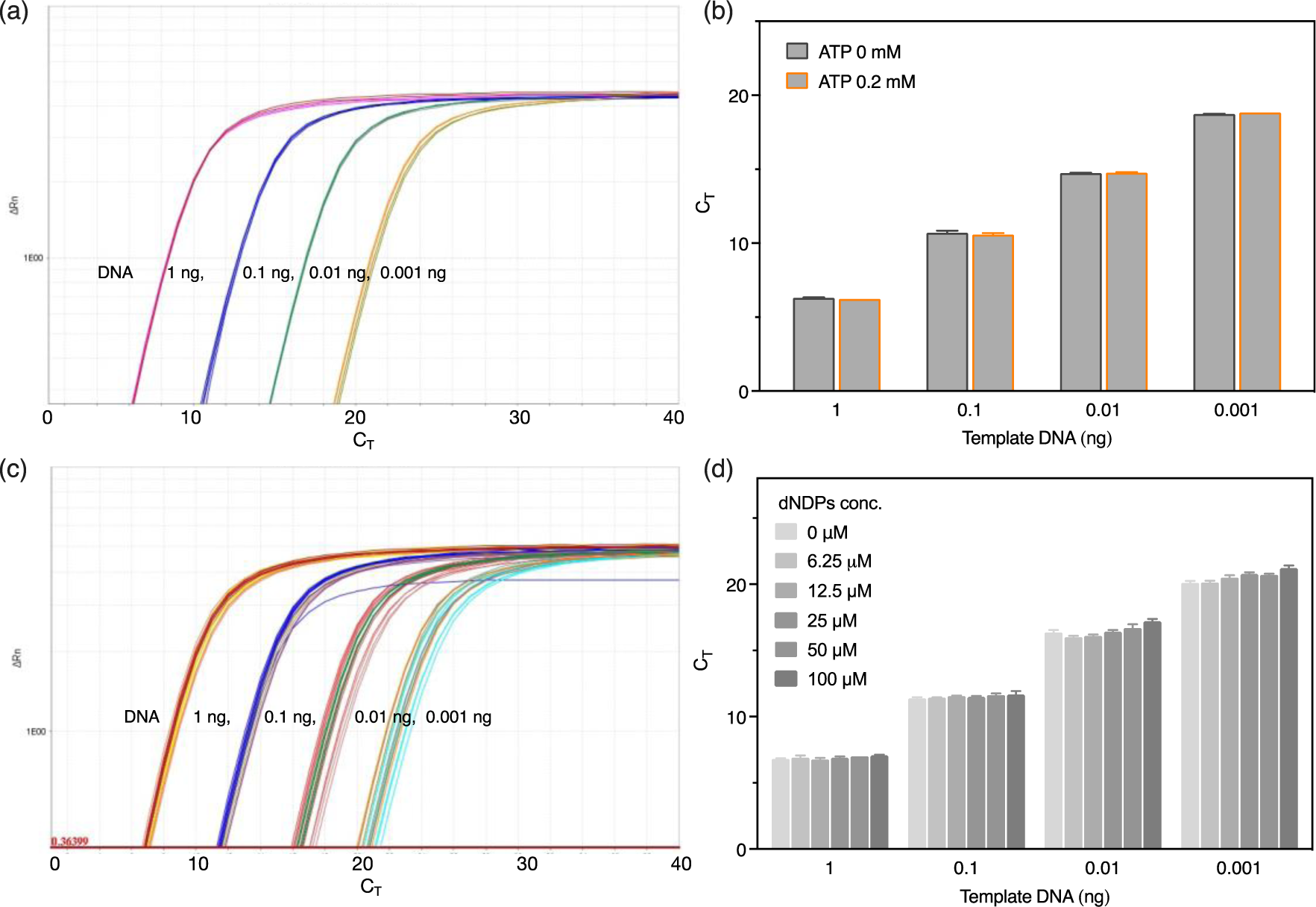
Amplification plots (a) and C_T_ values (b) of qPCR kits with and without 0.2 mM ATP added. Amplification plots (c) and C_T_ values (d) of qPCR kits with 0-100 µM dNDPs added.

